# Impact of trait measurement error on quantitative genetic analysis of computer vision derived traits

**DOI:** 10.1101/2025.06.02.657462

**Authors:** Ye Bi, Yijian Huang, Haipeng Yu, Gota Morota

**Author notes:** Corresponding author (GM). Email addresses (YB), (YH), (HY).

## Abstract

**Background:** Quantitative genetic analysis of image- or video-derived phenotypes is increasingly being performed for a wide range of traits. Pig body weight values estimated by a conventional approach or a computer vision system can be considered as two different measurements of the same trait, but with different sources of phenotyping error. Previous studies have shown that trait measurement error, defined as the difference between manually collected phenotypes and image-derived phenotypes, can be influenced by genetics, suggesting that the error is systematic rather than random and is more likely to lead to misleading quantitative genetic analysis results. Therefore, we investigated the effect of trait measurement error on genetic analysis of pig body weight (BW).

**Results:** Calibrated scale-based and image-based BW showed high coefficients of determination and goodness of fit. Genomic heritability estimates for scale-based and image-based BW were mostly identical across growth periods. Genomic heritability estimates for trait measurement error were consistently negligible, regardless of the choice of computer vision algorithm. In addition, genome-wide association analysis revealed no overlap between the top markers identified for scale-based BW and those associated with trait measurement error. Overall, the deep learning-based regressions outperformed the adaptive thresholding segmentation methods.

**Conclusions:** This study showed that manually measured scale-based and image-based BW phenotypes yielded the same quantitative genetic results. We found no evidence that BW trait measurement error could be influenced, at least in part, by genetic factors. This suggests that trait measurement error in pig BW does not contain systematic errors that could bias downstream genetic analysis.

## Background

Power of quantitative genetic analysis largely depends on quality and quantity of available phenotypes. Computer vision approaches are rapidly being developed to generate phenotypes for genetic analysis in livestock (Pérez-Enciso and Steibel, 2021). Computer vision coupled with cameras is one of the available precision livestock technologies to realize precision livestock farming, which aims to automate phenotyping or monitoring animals at the individual level on a continuous basis (Morota et al., 2018). An advantage of computer vision systems is that they can be installed in a barn with minimal disruption to daily farm operations and are relatively inexpensive, requiring only one or a few off-the-shelf consumer cameras (Kadlec et al., 2022). Pedigree-based or genomic-based genetic analysis of image- or video-derived traits is increasingly being performed in growth or morphological traits (Nye et al., 2020; Gorssen et al., 2022; Ricard et al., 2023; Manzanilla-Pech et al., 2023; Qin et al., 2024), carcass traits (Moore et al., 2017; Nakajima et al., 2018; Xie et al., 2021; Kaseja et al., 2024; Shen et al., 2024), behavior (Gorssen et al., 2022; Hollifield et al., 2024) and feed intake (Manzanilla-Pech et al., 2023). Computer vision is the core engine for phenotyping these live animal traits, except for carcass traits, which are collected after slaughter.

Computer vision is not necessarily always required to collect the above phenotypes, but traditional approaches often require additional hours of manual workload or the purchase of expensive equipment that requires careful maintenance. For example, collecting growth or morphological traits requires a digital scale with manual labor to restrain the animals or a costly walk-over weighing system. Similarly, manually monitoring individual feed intake requires constant observation and estimation of measurements by humans. Collecting behavioral traits requires many hours of behavioral video decoding by ethologists based on a formal ethogram. What distinguishes computer vision-based phenotyping from conventional phenotyping is the ability of the former to scale up and perform high-throughput phenotyping by automating all or part of the phenotyping procedures.

Consider a scenario where we are interested in phenotyping a specific target trait. This target trait, collected using conventional and image or video-based approaches, can be considered as two different measurements of the same character because they have different phenotyping errors. In other words, two measurements of the target trait are controlled by the same set of causal loci, but the source of phenotyping error is different. Then we can calculate a trait measurement error, which is defined as the difference between the conventional phenotypes and the image-derived phenotypes. Because these two measurements of the same trait have the same genetic architecture, it is assumed that trait measurement errors are random and nongenetic factors (Figure 1). However, some previous research has shown that trait measurement error can be influenced, at least in part, by genetic factors. For example, maize biomass was quantified using manual measurements and image-based estimates (Liang et al., 2018). Although they obtained the Pearson correlation coefficient of 0.95 between the manual measurements and image-based estimates, 58% of the trait measurement error in biomass was explained by genetic variation. Similarly, 27%-59% of the trait measurement error variance was controlled by genetic variation using single nucleotide polymorphism (SNP) data in sorghum height, even the Pearson correlation coefficient between manual measurements and image-based estimates was 0.96 (Zhou et al., 2021). They also identified a few significant genomic regions when trait measurement errors were used as phenotypes in genome-wide association studies (GWAS). These results suggest that phenotypes of some individuals are systematically underestimated and phenotypes of some other individuals are systematically overestimated. Because trait measurement error is systematic rather than random in these cases, if these image-derived phenotypes were used in quantitative genetic analysis, it could lead to misleading results.

**Figure 1:**
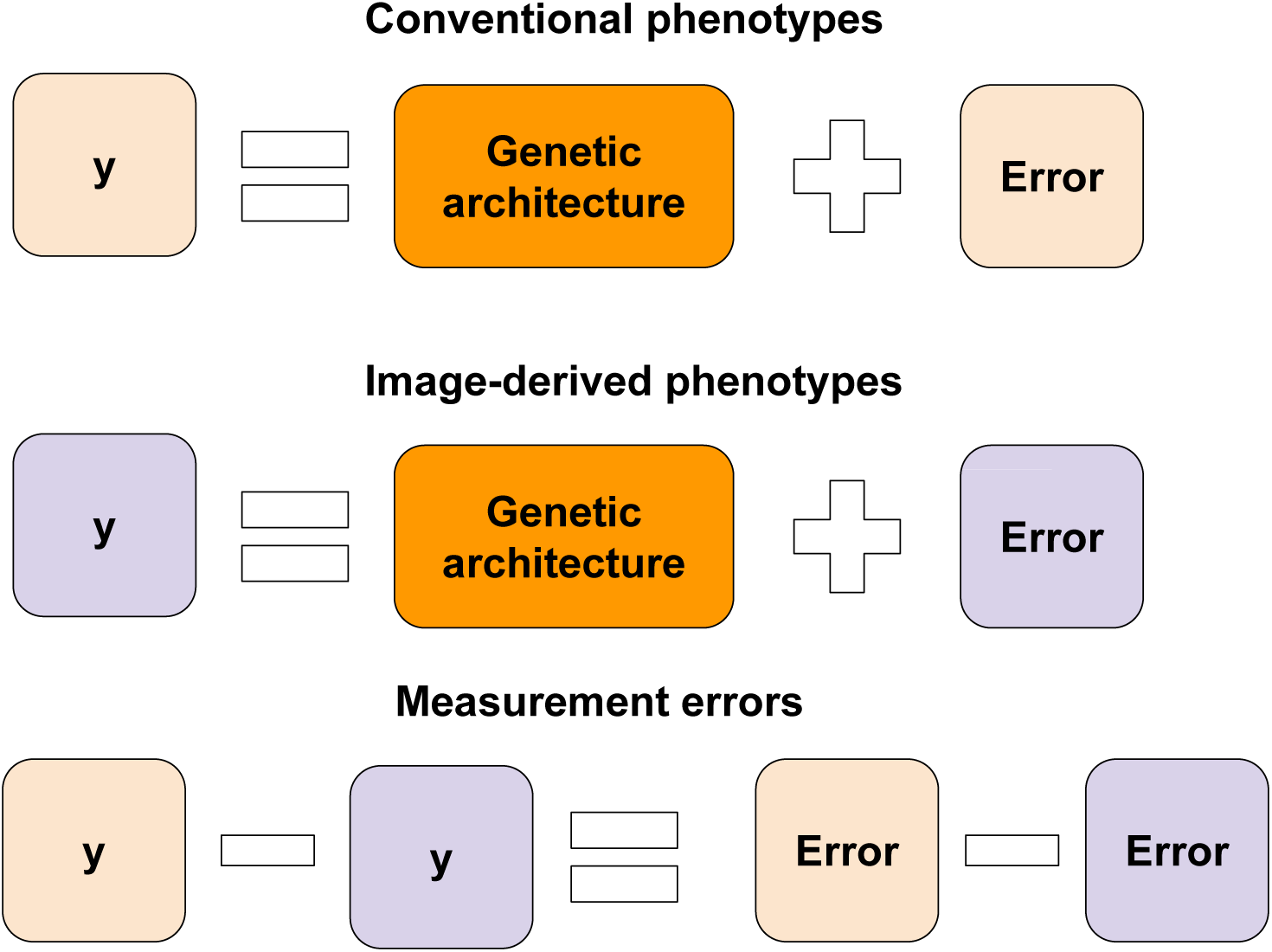
Concept of trait measurement error when using computer vision to generate image-derived phenotypes.

This type of problem can also occur in livestock. For example, the live body weight (BW) of animals is often estimated from a single top-view depth sensor camera (Yu et al., 2021; Wang et al., 2024). Height and volume information that can be estimated from a depth sensor camera is known to be highly correlated with scale-based BW in animals (Bi et al., 2023, 2025). When estimating volumetric measures from a depth sensor camera, it is assumed that the distance from the ground to the ventral abdomen is constant for all animals. However, this assumption is not valid if some sires or boars carry genetics that increase the distance from the ground to the ventral abdomen in their offspring, resulting in a systematic error in image-derived BW estimates. Although an increasing number of animal phenotypes are generated by computer vision, the effect of trait measurement error on quantitative genetic analysis has not been investigated. In addition, it is unknown how the choice of computer vision algorithms can influence trait measurement error in downstream genetic analysis.

Therefore, in this study, we investigated the impact of trait measurement error on the genetic analysis of pig BW. Specifically, the objectives of this study were 1) to estimate trait measurement error from manually collected and image-derived pig BW, 2) to evaluate how the choice of computer vision algorithms affects the genetic analysis of trait measurement error, and 3) to estimate genomic heritability and marker effects for trait measurement error.

## Materials and Methods

### Animals and data collection design

A subset of pig data analyzed in Bi et al. (2024, 2025) was used in this study. In short, we analyzed data from more than 600 crossbred pigs raised on a farm in North Carolina, USA. The pigs were part of a commercial finishing operation at Smithfield and were regularly weighed during four farm visits or time points (T1 to T4) between July and September 2023. The number of pigs phenotyped varied at each visit. Animals were fed a common diet and had unrestricted access to water. Pigs were weighed individually on a digital walk-in calibrated scale to ensure they were restrained and properly positioned for accurate measurements. Their BW ranged from 25 to 154 kg.

### Video acquisition setup

A depth video acquisition setup is detailed in our previous work (Bi et al., 2023, 2024, 2025). Briefly, an Intel RealSense D435 camera (Intel, Santa Clara, CA, USA) connected via USB 3.1 was strategically positioned 1.40 meters above a one-way exit lane equipped with a stand-on weighing system to capture top-view video. The camera’s dual stereo image sensors, with a horizontal field of view of 87^◦^ and a vertical field of view of 58^◦^, enabled depth estimation. Individual pigs were manually restrained to facilitate the collection of 10-12 second video segments at a frame rate of 30 frames per second and a resolution of 1280 × 720. While the pigs exhibited general stability in their body movements, their head and tail exhibited a comparatively dynamic range of motion. Simultaneously with the video recordings, the pigs’ BW was measured using a weight scale and individual identification was established by ear tag recognition.

### Depth data processing and quality control

The raw depth data captured in .bag video format was converted to PNG depth images and CSV depth map files using the *rs-convert* tool (Intel RealSense, 2023). Within each depth image, different colors represent different distances from the camera to the object, providing a visual representation of spatial depth. Each image was accompanied by a CSV file of identical dimensions, providing distance measurements in meters for each pixel.

To ensure the quality of the videos collected in the commercial environment for subsequent image analysis, we applied a customized YOLOv8s model (Jocher et al., 2023) during the preprocessing stage. We first annotated 100 depth images using Roboflow (Lin et al., 2022), labeling two categories: ”pig” (representing the main body of the pig) and ”block” (representing occlusions caused by farm technicians obstructing the camera view). After training the YOLOv8s model on these annotations, we detected objects within the depth images. Images were excluded if the block detection rate was greater than 0.1, the pig detection rate was less than 0.5, or no pig was detected. After preprocessing, the final dataset contained 20,356, 45,814, 44,525, and 23,968 depth images for T1 to T4, respectively.

### Adaptive thresholding segmentation

The first computer vision approach to derive image-based BW from depth images was adaptive thresholding segmentation followed by a regression model, as developed in a previous study for dairy cattle (Bi et al., 2023) and recently applied to. This approach extracts the biometric measurements of pigs and uses them to build regression models for BW prediction.

The OpenCV library (version 4.7.0) was used in Python (version 3.8.16) for the thresholding method. First, the depth images were cropped to eliminate extraneous surroundings. In hue, saturation, and value (HSV) color mode, the hue channel was extracted as a threshold for converting the images to binary format, where hues above the threshold were converted to black and those below were converted to white. An adaptive range from 15 degrees below to 5 degrees above the mean of the hue was applied. In general, lower thresholds were preferred because they captured more of the animal’s body. To determine the optimal threshold, we used the OpenCV function *boundingRect* to draw a bounding box around the largest contour. If the four corners of the bounding box were less than five pixels from the edge, the hue threshold was increased by one unit until the four corners were more than five pixels from the edge of the image. The OpenCV function *findcontour* was then used to find the contour with the largest area. Finally, the OpenCV function *minAreaRect* was used to draw the minimum bounding rectangle of the pig’s body contour. Due to the significant movement of the pig’s head and tail during the video data acquisition, we excluded these areas from the depth images. The ratio of body width to body length was used to remove the neck and tail horizontally and vertically within each image. Finally, the contour was identified and the bounding box was drawn on the binary image. The bounding box and depth information associated with the identified contour were retained for BW regression analysis.

Four biometric features were extracted from each image: dorsal length, abdominal width, height, and volume. Dorsal length and abdominal width were estimated by drawing the minimum area bounding box around the pig’s body using the OpenCV function *minAreaRect*. Height was determined using an average-based approach where missing data (zeros) within the contour pixels in the depth CSV file were replaced with the average distance. The average height of the pig was calculated as 1.40 meters minus the average distance from the camera to the pig contour. Volume was calculated by summing the height values of pixels within the pig contour. After all images were processed, the medians of dorsal length, abdominal width, height, and volume were calculated to produce a single final estimate for each video. These image-derived features were then used as predictors to build regression models for predicting pig body weight. Ordinary least squares (OLS) and random forest (RF) models were applied using the caret R package (Kuhn, 2015). To improve the performance of the RF model and reduce the risk of overfitting, 5-fold cross-validation was performed within the training dataset.

### Deep learning models

The second computer vision approach used to estimate BW was deep learning models. The main idea is to directly predict pig BW by feeding the entire depth image to neural network models as shown in recent studies (Bi et al., 2024, 2025). Deep learning models integrate feature extraction and regression into a single-step process. A brief description is given below, as details are given in a recent study (Bi et al., 2025).

We first converted depth CSV files containing distance information in meters to depth grayscale images to ensure consistent distance representation. This conversion was done using the *pyplot* function in the matplotlib library (version 3.7.1) in Python. The grayscale images, originally 1270 × 720 pixels in size, were resized and padded to a square dimension of 150 × 150. We evaluated the feature extraction capabilities of two deep learning model families using transfer learning: MobileViT and ResNet50, both pre-trained on ImageNet. Modifications included removing the classifier layer, retaining the backbone, adding a 50% dropout layer, and incorporating a single-node dense layer with linear activation and L2 regularization for BW prediction. The main architectures are summarized below.

MobileViT combines the strengths of convolutional neural networks and transformers by integrating spatial inductive biases with global processing capabilities (Mehta and Rastegari, 2021). It uses a hybrid block that applies standard and pointwise convolutions to capture local features, transforms these into non-overlapping patches for long-range dependency modeling via transformers, and then re-folds and fuses features with the input tensor. We evaluated the small MobileViT architecture, MobileViT-S. ResNet50 was used as the baseline model. It uses residual blocks and applies an identity shortcut connection to skip one or more layers to mitigate the degradation problem, allowing for easier optimization and increased accuracy with depth (He et al., 2016). MobileViT is expected to provide greater efficiency than ResNet50 by leveraging a hybrid architecture that integrates convolutional layers for local feature extraction with transformer layers for capturing long-range dependencies, resulting in a more parameter-efficient model. In contrast, ResNet50 relies on residual connections to improve optimization and accuracy with depth, but its architecture generally requires more computational resources as model depth increases.

The deep learning models were implemented using TensorFlow GPU 2.14.0 and Keras 2.14.0 in Python 3.9.0, running on Virginia Tech’s Advanced Research Computing cluster with two 80 GB NVIDIA DGX A100 GPUs. Models were trained on 150 × 150 images with a batch size of 100 for 300 epochs, using mean squared error as the loss function. The *ReduceLROnPlateau* callback dynamically reduced the learning rate by 0.1 when validation loss stagnated for 75 epochs, with a minimum threshold of 0.000001, while *ModelCheckpoint* saved the best model based on validation loss. Validation loss was the criterion for selecting the optimal model parameters.

### Goodness of fit metrics

The coefficient of determination (R^2^) and the mean absolute percentage error (MAPE) were used to evaluate the performance of the computer vision models.

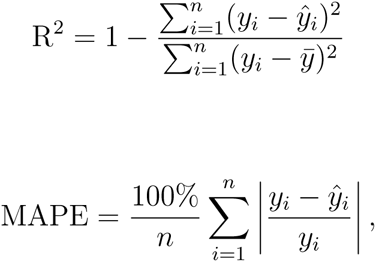

where *y_i_* is the scale-based BW of the *i*th animal, *ŷ_i_* is the image-based predicted BW of the *i*th animal, and *ȳ* is the average scale-based BW.

### Quantitative genetic modeling

All pigs were genotyped using 50K SNP chips. After quality control by removing markers with minor allele frequencies less than 0.05, a total of 46,407 to 46,424 SNPs remained for analysis depending on different growth periods and computer vision approaches used because number of available pigs differed.

Genomic heritability estimates of scale-based BW, image-based BW, and trait measurement error BW defined as their absolute differences were obtained using restricted maximum likelihood method (Patterson and Thompson, 1971) in the genomic best linear unbiased prediction (GBLUP) model (VanRaden, 2008). The GBLUP model used was

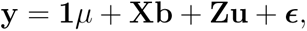

where **y** is a vector of phenotypes; **1** is the vector of ones; *µ* is the overall mean; **X** is the incidence matrix of fixed effects; **b** is a vector of fixed effects; **Z** is the incidence matrix relating animals to their phenotypic records; and **u** is a vector of the random additive genetic values of animals; and ***ɛ*** ∼ N(**0**, **I***σ_ɛ_*^2^), is a vector of residuals. We assumed **u** ∼ N(**0**, **G***σ*_u_^2^), where 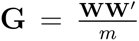*^′^* is the genomic relationship matrix; *σ*^2^ is the additive genetic variance;**W** is a centered and standardized SNP marker matrix; and *m* is the total number of SNP markers. The fixed effects included, sex, birth farm, pen density, age, and parity of dam.

We used the following single-marker GWAS (Kennedy et al., 1992; Yu et al., 2006) for scale-based BW, image-based BW, and their trait measurement errors.

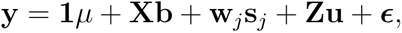

where **w***_j_* is the *j*th SNP marker to be tested and **s***_j_* is a vector of fixed *j*th SNP effect. The remaining terms are identical to those in the GBLUP model. All models were fitted using the rrBLUP R package (Endelman, 2011). Since our main interest is whether we can correctly detect SNP markers with large effects, we evaluated whether the top 10 markers detected in scale-based BW were among the top 100 markers in image-based BW or trait measurement error BW using -log 10 p-values. We hypothesized that among a total of SNPs, almost all of the top 10 markers detected in scale-based BW would also be correctly detected in the top 100 markers of image-based BW, whereas none of the top 10 markers detected in scale-based BW would be detected in the top 100 markers of trait measurement error BW. We focused on SNP markers with top GWAS peaks because it is typically not of interest to correctly identify the ranking of SNPs with small effects. All genetic analyses were performed at each growth period to investigate whether there are specific time points at which trait measurement error is genetically influenced.

## Results

Phenomic analysis with deep regression resulted in more pigs with phenotypes for genetic analysis than thresholding segmentation (Table 1). This is because the adaptive thresholding approach was unable to process some pigs with noisy image data, whereas deep learning-based regression was robust to variable image quality and was able to process noisy image data. The amount of trait measurement error varied between pigs at different time points. Figure 2 shows pigs that had the highest trait measurement errors at at least two time points and lowest trait measurement errors at least two time points. The results indicate that some pigs had consistently high trait measurement errors over time, while other pigs had consistently low trait measurement errors.

**Figure 2:**
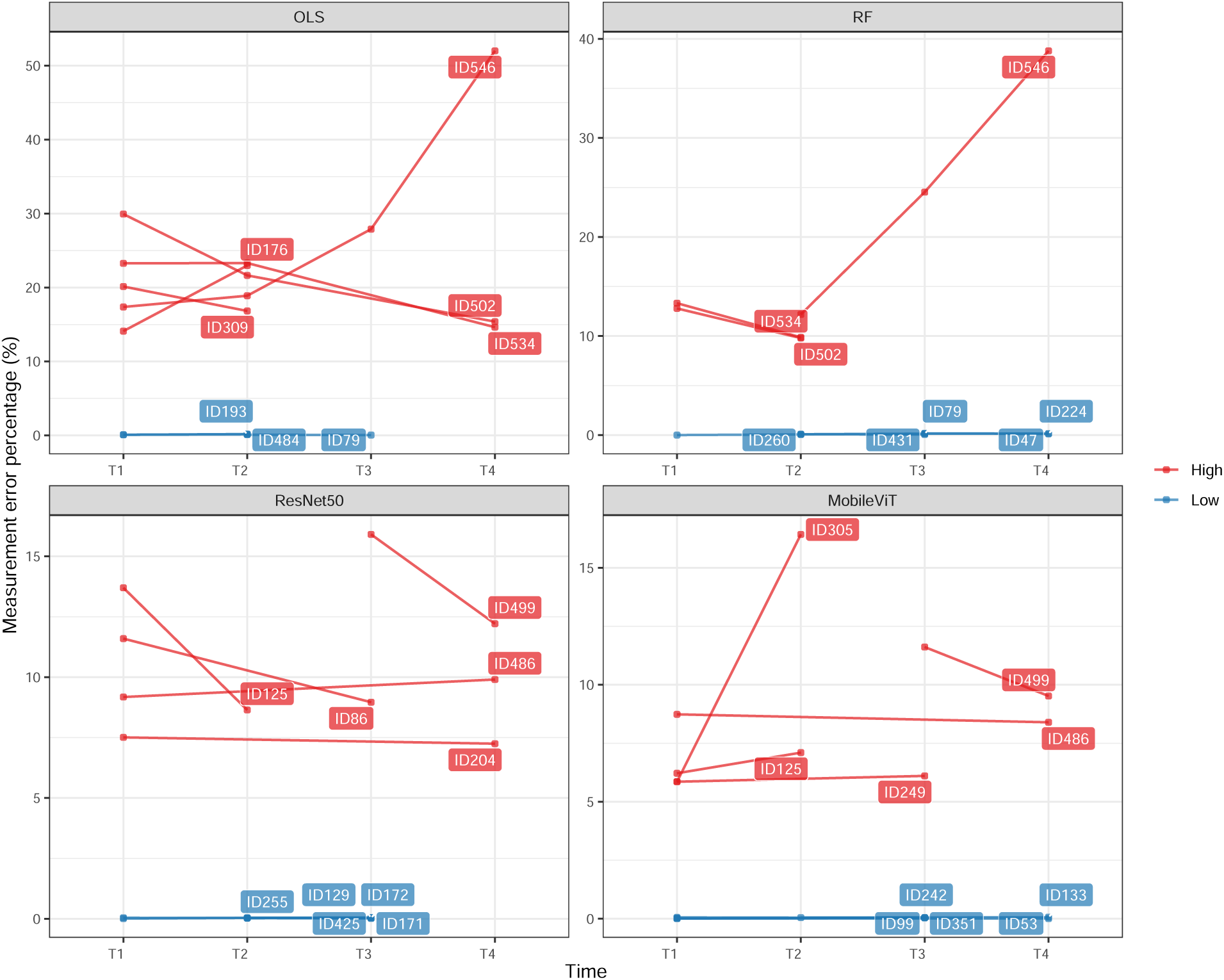
Illustration of pigs that had the highest trait measurement errors on at least two time points and the lowest trait measurement errors on at least two time points.

**Table 1:**
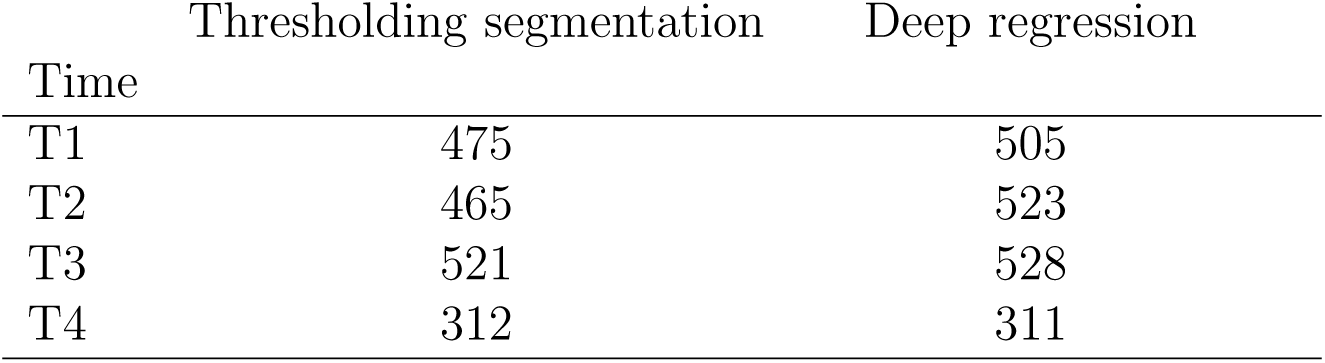
Number of pigs with image-derived phenotypes produced by two computer vision appraoches. T1 to T4 denote four farm visits.

Image-derived BW were estimated by training models on video data captured by a depth camera. A total of four image analysis models were used, including adaptive thresholding segmentation coupled with either OLS or RF, and deep regression based on ResNet50 and MobileViT. Coefficients of determination for image-derived and scale-based body BW were at least 0.95 for RF-based adaptive thresholding segmentation approaches and deep regression across different time points (Table 2), showing strong agreement. Similarly, the MAPE was always less than 2.5% for the RF-based adaptive thresholding segmentation approaches and the deep regressions. Adaptive thresholding segmentation with OLS resulted in consistently lower R^2^ and higher MAPE than the other approaches. T3, which had the largest number of pigs available, showed the best performance for deep regression. Trait measurement errors expressed as percentages showed that deep regression provided the lowest errors across all growth periods, followed by RF- and OLS-based adaptive thresholding segmentation (Figure 3). The mean trait measurement error percentages were 4.57%, 2.13%, 1.64%, and 1.61% for OLS-based adaptive thresholding segmentation, RF-based adaptive thresholding segmentation, ResNet50, and MobileViT, respectively.

**Figure 3:**
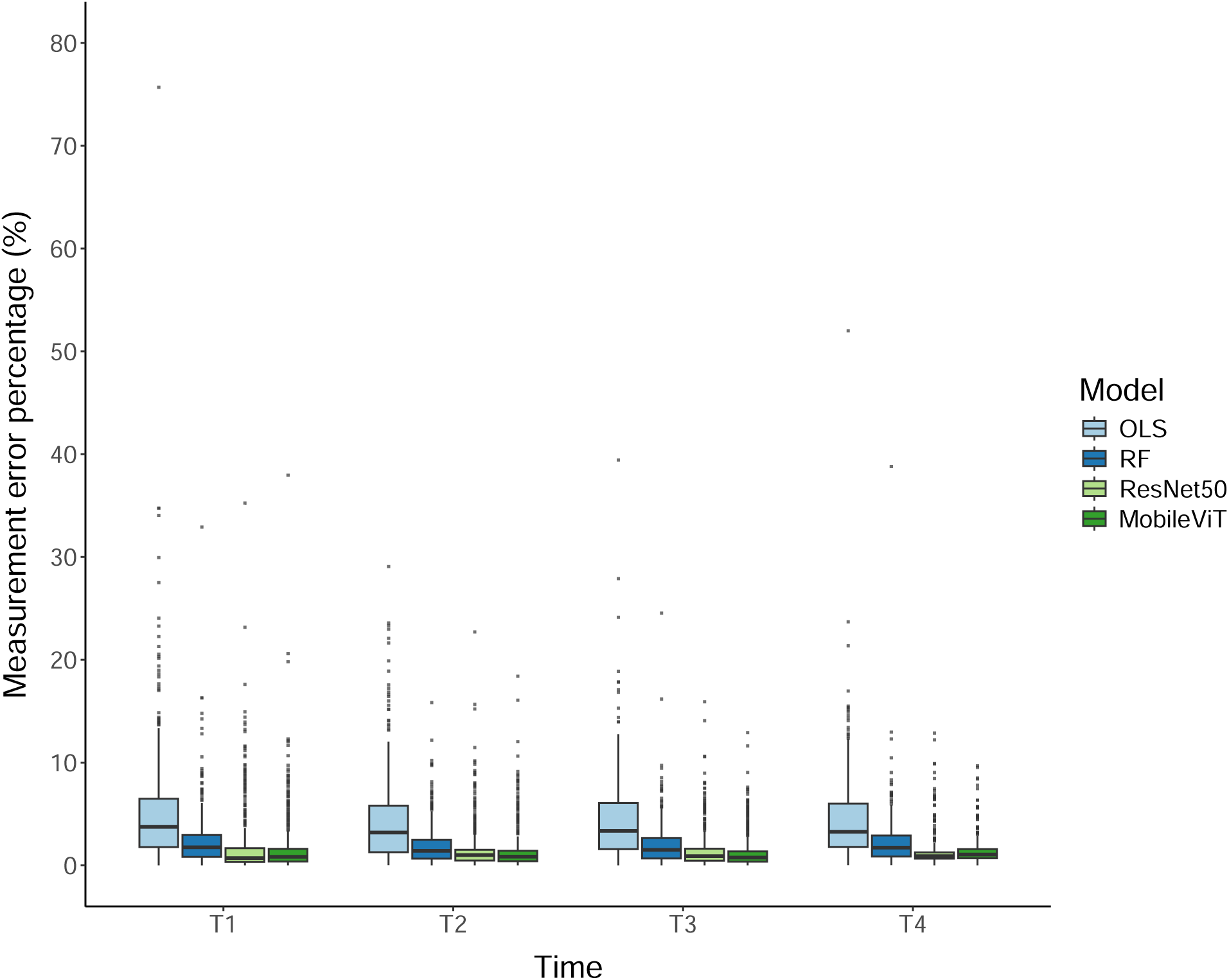
Trait measurement error percentages of four computer vision approaches across four data collection points. OLS: ordinarry least squares. RF: Random forest.

**Table 2:** Coefficient of determination and mean absolute percentage error between scale-based body weight and image-derived body weight in the pig data. T1 to T4 denote four farm visits.

| Time | Thresholding segmentation |  |  |  | Deep regression |  |  |  |
| --- | --- | --- | --- | --- | --- | --- | --- | --- |
|  | OLS<sup>1</sup> |  | RF<sup>2</sup> |  | ResNet50 |  | MobileViT |  |
|  | R<sup>2</sup><sup>3</sup> | MAPE<sup>4</sup> | R<sup>2</sup> | MAPE | R<sup>2</sup> | MAPE | R<sup>2</sup> | MAPE |
| T1 | 0.72 | 5.33 | 0.95 | 2.45 | 0.93 | 1.87 | 0.94 | 1.68 |
| T2 | 0.84 | 4.29 | 0.97 | 1.95 | 0.95 | 1.54 | 0.96 | 1.44 |
| T3 | 0.82 | 4.24 | 0.96 | 1.96 | 0.97 | 1.44 | 0.98 | 1.27 |
| T4 | 0.80 | 4.57 | 0.95 | 2.30 | 0.96 | 1.44 | 0.97 | 1.45 |

<sup>1</sup> Ordinary least squares
<sup>2</sup> Random forests
<sup>3</sup> Coefficient of determination
<sup>4</sup> Mean absolute percentage error

Genomic heritability estimates for scale-based BW ranged from 0.21 to 0.55 across time points and numbers of pigs available (Table 3). Genomic heritability estimates for manual and image-based BW were mostly identical across time points, indicating that manually measured and image-based phenotypes yield the same genetic inference results. Agreement was particularly evident for the deep regression approaches, with differences in genomic heritability estimates ranging from 0 to 0.03. On the other hand, genomic heritability estimates for BW obtained from the OLS-based adaptive segmentation approach differed most from those of scale-based BW. Genomic heritability estimates for measurement error were consistently close to zero and negligible.

**Table 3:** Genomic heritability estimates for scale-based body weight, image-based body weight, and trait measurement error.

| Time | $h^2$ type<sup>1</sup> | Thresholding | | Deep regression | |
| --- | --- | --- | --- | --- | --- |
|  |  | OLS<sup>2</sup> | RF<sup>3</sup> | ResNet50 | MobileViT |
| T1 | $h^2_{\text{BW}^4}$ | 0.54 | | 0.55 | |
| | $h^2_{\text{CV}^5}$ | 0.44 | 0.56 | 0.55 | 0.55 |
| | $h^2_{\text{TME}^6}$ | 0.00 | 0.02 | 0.03 | 0.03 |
| T2 | $h^2_{\text{BW}}$ | 0.46 | | 0.39 | |
| | $h^2_{\text{CV}}$ | 0.33 | 0.40 | 0.39 | 0.40 |
| | $h^2_{\text{TME}}$ | 0.00 | 0.00 | 0.00 | 0.00 |
| T3 | $h^2_{\text{BW}}$ | 0.31 | | 0.29 | |
| | $h^2_{\text{CV}}$ | 0.36 | 0.32 | 0.31 | 0.32 |
| | $h^2_{\text{TME}}$ | 0.02 | 0.00 | 0.00 | 0.00 |
| T4 | $h^2_{\text{BW}}$ | 0.23 | | 0.21 | |
| | $h^2_{\text{CV}}$ | 0.27 | 0.25 | 0.21 | 0.20 |
| | $h^2_{\text{TME}}$ | 0.00 | 0.03 | 0.00 | 0.00 |

<sup>1</sup> Types of genomic heritability estimate
<sup>2</sup> Ordinary least squares
<sup>3</sup> Random forests
<sup>4</sup> Genomic heritability estimate for scale-based body weight
<sup>5</sup> Genomic heritability estimate for predicted weight using computer vision
<sup>6</sup> Genomic heritability estimate for trait measurement error

Table 4 shows the number of top 10 markers detected by scale-based BW that are correctly detected in the top 100 markers of image-based BW, and the number of top 10 markers detected by scale-based BW that are incorrectly detected in the top 100 markers of trait measurement error BW in GWAS. If computer vision approaches can accurately estimate BW without systematic error, the top 10 markers identified by scale-based methods should also be among the top 100 markers identified by image-based methods. However, these top 10 scale-based markers would not be found among the top 100 markers derived from trait measurement errors. The GWAS analysis showed that both the RF-based adaptive thresholding segmentation and the deep regression approaches can always correctly identify the top 10 markers from scale-based BW within their top 100 markers. However, the OLS-based adaptive thresholding approach correctly identified only 4 to 7 top 10 markers. When GWAS was performed on trait measurement errors, none of the top 10 markers from scale-based BW were detected by the computer vision approaches within their top 100 markers.

**Table 4:** Number of top 10 markers detected in scale-based body weight that is correctly detected in the top 100 markers of image-based body weight (Agreement) and number of the top 10 markers detected in scale-based body weight that is incorrectly detected in the top 100 markers of trait measurement error body weight (TME) in the genome-wide association studies. T1 to T4 denote four farm visits.

| Time | Thresholding segmentation |  |  |  | Deep regression |  |  |  |
| --- | --- | --- | --- | --- | --- | --- | --- | --- |
|  | OLS<sup>1</sup> |  | RF<sup>2</sup> |  | ResNet50 |  | MobileViT |  |
|  | Agreement | TME | Agreement | TME | Agreement | TME | Agreement | TME |
| T1 | 4/10 | 0/10 | 10/10 | 0/10 | 10/10 | 0/10 | 10/10 | 0/10 |
| T2 | 7/10 | 0/10 | 10/10 | 0/10 | 10/10 | 0/10 | 10/10 | 0/10 |
| T3 | 7/10 | 0/10 | 10/10 | 0/10 | 10/10 | 0/10 | 10/10 | 0/10 |
| T4 | 4/10 | 0/10 | 10/10 | 0/10 | 10/10 | 0/10 | 10/10 | 0/10 |

<sup>1</sup> Ordinary least squares
<sup>2</sup> Random forests

## Discussion

Computer vision approaches are rapidly being developed to accelerate phenomic research for livestock genetic analysis. However, more work is needed before the incorporation of image-based phenotypes into genetic evaluation analysis can be considered. There is a possibility that the reliability of phenotyping may decrease as phenotyping becomes increasingly large-scale in noisy heterogeneous environments, and systematic errors that may arise for specific animals as a result of phenotyping may be a serious concern.

This study investigated the influence of trait measurement error on downstream genetic analysis of BW in pigs phenotyped under commercial environmental conditions. Continuous recording of BW is becoming increasingly important because BW can be used to derive residual feed intake, which is viewed as a proxy for difficult to measure feed efficiency (Patience et al., 2015). Here, trait measurement error was defined as the difference between scaled and computer vision derived BW. Previous studies have suggested that downstream quantitative genetic analysis may be compromised even in a scenario where conventionally measured phenotypes and computer vision-derived phenotypes are highly correlated or have high goodness of fit. Although ensuring the safety of computer vision-derived phenotypes for use in downstream animal quantitative genetics is an important first step in data analysis, it has not received much attention in the literature. One possible reason for the lack of this type of study is that it requires phenotyping animals using both conventional and computer vision approaches, which requires intensive manual work and image analysis. Therefore, in the current study, we collected both scale-based and image-based BW and evaluated the consequences of using image-based BW as phenotypes in quantitative genetic analysis by deriving trait measurement error.

We found that some pigs had consistently high measurement errors across different time points, suggesting that these pigs were always difficult to accurately predict BW from images (Figure 2). In addition, OLS-based adaptive thresholding performed least well than competing approaches in inference tasks. For example, the OLS-based adaptive thresh-olding approach showed relatively low R^2^ and high MAPE between scale-based BW and image-based BW, image-based genomic heritability estimates sometimes differed by 0.1 from scale-based estimates, and only 4 to 7 of the top 10 GWAS markers detected in scale-based BW were correctly detected in the top 100 GWAS markers of image-based BW. However, these concerns did not lead to evidence of a genetic component to trait measurement error. Genomic heritability estimates for trait measurement error were always zero or negligible across growth periods, regardless of the choice of the computer vision approaches (Table 3). Also, none of the top 10 GWAS markers detected in scale-based BW were detected in the top 100 GWAS markers of trait measurement error. Our GWAS results are in line with a previous work reporting no significant difference in GWAS peaks between simulated manual and image-based maize tassel characters (Gage et al., 2018). Thus, it is likely that trait measurement error in image-based pig BW does not contain systematic errors characterized by genetics that could potentially bias downstream genetic analyses. A previous study reported that image-based phenotypes tended to overestimate pedigree-based heritability estimates relative to manually collected phenotypes for sheep growth traits (Qin et al., 2024), but we did not find the similar trend in our study.

Taken together, our results suggest that manual scale-based and image-based BW are controlled by the same set of causal loci, and their difference in phenotypic values is due to different phenotyping errors. Thus, the trait measurement error of BW during pig growth is generated by phenotyping errors that are not genetic, regardless of the computer vision approaches used. In particular, OLS-based adaptive thresholding did not produce BW as accurately as other competing approaches, but its trait measurement error was still free of genetic factors. It is worth noting that although manually measured phenotypes are often considered ground truth in computer vision analysis, even manual measurements have some degree of phenotyping error due to human or mechanical error. As the phenotyping error due to computer vision becomes smaller, the accuracy of phenotypes is expected to improve further.

## Conclusions

The advent of precision livestock farming offers increasing opportunities to use new technologies to generate phenotypes. Our results provide valuable insights into the effect of trait measurement error on quantitative genetic analysis in pigs when using computer vision technology. Our study found no evidence that trait measurement error includes genetic factors during pig growth. It is likely that image-derived BW in pigs can be used without reservation for downstream quantitative genetic analysis.

## Author contribution statement

Ye Bi: Formal analysis, Investigation; Data curation; Visualization; Writing-original draft; Writing-review & editing. Yijian Huang: Investigation; Data curation; Project administration; Writing-review & editing. Haipeng Yu: Methodology; Writing-review & editing. Gota Morota: Conceptualization; Methodology; Formal analysis; Investigation; Data curation; Funding acquisition; Project administration; Supervision; Writing-original draft; Writing-review & editing.

## Acknowledgments

The authors acknowledge Advanced Research Computing at Virginia Tech (https://arc.vt.edu/) for providing computational resources and technical support that have contributed to the results reported within this paper.

## Data availability

The data supporting the results of this article are the property of Smithfield Premium Genetics, and this information is commercially sensitive and therefore cannot be made available.

## Ethics declarations

### Ethics approval and consent to participate

All biological material used in this study was collected as part of routine data collection at Smithfield Premium Genetics and not specifically for the purpose of this project. Therefore, ethics committee approval was not required.

### Consent for publication

Not applicable.

## References

1. Bi, Y., Campos, L. M., Wang, J., Yu, H., Hanigan, M. D., and Morota, G. (2023). Depth video data-enabled predictions of longitudinal dairy cow body weight using thresholding and Mask R-CNN algorithms. Smart Agricultural Technology, 6:100352.

2. Bi, Y., Huang, Y., Xuan, J., and Morota, G. (2025). Industry-scale prediction of video-derived pig body weight using efficient convolutional neural networks and vision transformers. Biosystems Engineering, page In press.

3. Bi, Y., Xuan, J., Huang, Y., and Morota, G. (2024). 465 Comparative analysis of semantic segmentation and deep regression models with supervised pre-training for accurate prediction of pig body weight from video data: Insights from industry-scale datasets. Journal of Animal Science, 102(Supplement 3):414–415.

4. Endelman, J. B. (2011). Ridge regression and other kernels for genomic selection with R package rrblup. The plant genome, 4(3).

5. Gage, J. L., de Leon, N., and Clayton, M. K. (2018). Comparing genome-wide association study results from different measurements of an underlying phenotype. G3: Genes, Genomes, Genetics, 8(11):3715–3722.

6. Gorssen, W., Winters, C., Meyermans, R., D’Hooge, R., Janssens, S., and Buys, N. (2022). Estimating genetics of body dimensions and activity levels in pigs using automated pose estimation. Scientific Reports, 12(1):15384.

7. He, K., Zhang, X., Ren, S., and Sun, J. (2016). Deep residual learning for image recognition. In Proceedings of the IEEE Conference on Computer Vision and Pattern Recognition, pages 770–778.

8. Hollifield, M. K., Chen, C.-Y., Psota, E., Holl, J., Lourenco, D., and Misztal, I. (2024). Estimating genetic parameters of digital behavior traits and their relationship with production traits in purebred pigs. Genetics Selection Evolution, 56(1):29.

9. Intel RealSense (Accessed: March 1, 2023.). Intel realsense github. https://github.com/IntelRealSense.

10. Jocher, G., Chaurasia, A., and Qiu, J. (2023). Ultralytics YOLO.

11. Kadlec, R., Indest, S., Castro, K., Waqar, S., Campos, L. M., Amorim, S. T., Bi, Y., Hanigan, M. D., and Morota, G. (2022). Automated acquisition of top-view dairy cow depth image data using an RGB-D sensor camera. Translational Animal Science, 6(4):txac163.

12. Kaseja, K., Lambe, N., Yates, J., Smith, E., and Conington, J. (2024). Genome wide association studies for carcass traits measured by video image analysis in crossbred lambs. Meat Science, 214:109518.

13. Kennedy, B., Quinton, M., and Van Arendonk, J. (1992). Estimation of effects of single genes on quantitative traits. Journal of Animal Science, 70(7):2000–2012.

14. Kuhn, M. (2015). Caret: classification and regression training. Astrophysics Source Code Library, pages ascl–1505.

15. Liang, Z., Pandey, P., Stoerger, V., Xu, Y., Qiu, Y., Ge, Y., and Schnable, J. C. (2018). Conventional and hyperspectral time-series imaging of maize lines widely used in field trials. Gigascience, 7(2):gix117.

16. Lin, Q., Ye, G., Wang, J., and Liu, H. (2022). Roboflow: a data-centric workflow management system for developing ai-enhanced robots. In Conference on Robot Learning, pages 1789–1794. PMLR.

17. Manzanilla-Pech, C. I., Stephansen, R. B., and Lassen, J. (2023). Genetic parameters for feed intake and body weight in dairy cattle using high-throughput 3-dimensional cameras in Danish commercial farms. Journal of Dairy Science, 106(12):9006–9015.

18. Mehta, S. and Rastegari, M. (2021). MobileViT: light-weight, general-purpose, and mobile-friendly vision transformer. arXiv preprint arXiv:2110.02178.

19. Moore, K., Mrode, R., and Coffey, M. (2017). Genetic parameters of visual image analysis primal cut carcass traits of commercial prime beef slaughter animals. Animal, 11(10):1653– 1659.

20. Morota, G., Ventura, R. V., Silva, F. F., Koyama, M., and Fernando, S. C. (2018). Big data analytics and precision animal agriculture symposium: Machine learning and data mining advance predictive big data analysis in precision animal agriculture. Journal of animal science, 96(4):1540–1550.

21. Nakajima, A., Kawaguchi, F., Uemoto, Y., Fukushima, M., Yoshida, E., Iwamoto, E., Akiyama, T., Kohama, N., Kobayashi, E., Honda, T., et al. (2018). A genome-wide association study for fat-related traits computed by image analysis in Japanese Black cattle. Animal Science Journal, 89(5):743–751.

22. Nye, J., Zingaretti, L. M., and Pérez-Enciso, M. (2020). Estimating conformational traits in dairy cattle with DeepAPS: a two-step deep learning automated phenotyping and segmentation approach. Frontiers in genetics, 11:513.

23. Patience, J. F., Rossoni-Serão, M. C., and Gutiérrez, N. A. (2015). A review of feed efficiency in swine: biology and application. Journal of animal science and biotechnology, 6:1–9.

24. Patterson, H. D. and Thompson, R. (1971). Recovery of inter-block information when block sizes are unequal. Biometrika, 58(3):545–554.

25. Pérez-Enciso, M. and Steibel, J. P. (2021). Phenomes: the current frontier in animal breeding. Genetics Selection Evolution, 53(1):22.

26. Qin, Q., Zhang, C., Liu, Z., Wang, Y., Kong, D., Zhao, D., Zhang, J., Lan, M., Wang, Z., Alatan, S., et al. (2024). Estimation of the genetic parameters of sheep growth traits based on machine vision acquisition. animal, page 101196.

27. Ricard, A., Crevier-Denoix, N., Pourcelot, P., Crichan, H., Sabbagh, M., Dumont-Saint-Priest, B., and Danvy, S. (2023). Genetic analysis of geometric morphometric 3D visuals of french jumping horses. Genetics Selection Evolution, 55(1):63.

28. Shen, Y., Chen, Y., Zhang, S., Wu, Z., Lu, X., Liu, W., Liu, B., and Zhou, X. (2024). Smartphone-based digital phenotyping for genome-wide association study of intramuscular fat traits in longissimus dorsi muscle of pigs. Animal Genetics, 55(2):230–237.

29. VanRaden, P. (2008). Efficient methods to compute genomic predictions. Journal of Dairy Science, 91(11):4414–4423.

30. Wang, J., Hu, Y., Xiang, L., Morota, G., Brooks, S. A., Wickens, C. L., Miller-Cushon, E. K., and Yu, H. (2024). ShinyAnimalCV: open-source cloud-based web application for object detection, segmentation, and three-dimensional visualization of animals using computer vision. Journal of Animal Science, 102:skad416.

31. Xie, L., Qin, J., Rao, L., Tang, X., Cui, D., Chen, L., Xu, W., Xiao, S., Zhang, Z., and Huang, L. (2021). Accurate prediction and genome-wide association analysis of digital intramuscular fat content in longissimus muscle of pigs. Animal Genetics, 52(5):633–644.

32. Yu, H., Lee, K., and Morota, G. (2021). Forecasting dynamic body weight of nonrestrained pigs from images using an RGB-D sensor camera. Translational Animal Science, 5(1):txab006.

33. Yu, J., Pressoir, G., Briggs, W. H., Bi, I. V., Yamasaki, M., Doebley, J. F., McMullen, M. D., Gaut, B. S., Nielsen, D. M., Holland, J. B., et al. (2006). A unified mixed-model method for association mapping that accounts for multiple levels of relatedness. Nature Genetics, 38(2):203.

34. Zhou, Y., Kusmec, A., Mirnezami, S. V., Attigala, L., Srinivasan, S., Jubery, T. Z., Schnable, J. C., Salas-Fernandez, M. G., Ganapathysubramanian, B., and Schnable, P. S. (2021). Identification and utilization of genetic determinants of trait measurement errors in image-based, high-throughput phenotyping. The Plant Cell, 33(8):2562–2582.

